# The ExAC Browser: Displaying reference data information from over 60,000 exomes

**DOI:** 10.1101/070581

**Authors:** Konrad J. Karczewski, Ben Weisburd, Brett Thomas, Douglas M. Ruderfer, David Kavanagh, Tymor Hamamsy, Monkol Lek, Kaitlin E. Samocha, Beryl B. Cummings, Daniel Birnbaum, The Exome Aggregation Consortium, Mark J. Daly, Daniel G. MacArthur

## Abstract

Worldwide, hundreds of thousands of humans have had their genomes or exomes sequenced, and access to the resulting data sets can provide valuable information for variant interpretation and understanding gene function. Here, we present a lightweight, flexible browser framework to display large population datasets of genetic variation. We demonstrate its use for exome sequence data from 60,706 individuals in the Exome Aggregation Consortium (ExAC). The ExAC browser provides gene- and transcript-centric displays of variation, a critical view for clinical applications. Additionally, we provide a variant display, which includes population frequency and functional annotation data as well as short read support for the called variant. This browser is open-source, freely available, and has already been used extensively by clinical laboratories worldwide.

## Introduction

Recently, large reference datasets, such as those from the 1000 Genomes Project Consortium^1^, Exome Sequencing Project (ESP)^2^, and Exome Aggregation Consortium (ExAC)^3^, have become publicly available for the benefit of the biomedical community. These projects release raw data in the form of variant call format (VCF) files, but these files require bioinformatics expertise to parse and synthesize. Genome browsers, such as the UCSC genome browser^4^, have become a popular method for non-technical audiences to visualize large genome-scale datasets.

Additionally, browsers of variation data, including the Exome Variant Server (EVS) from ESP and the 1000 Genomes Browser, have been developed to present population data, but these are limited in the data they display. For instance, deviations in coverage, which affect one's confidence of the absence of variation, are not natively shown: EVS contains a link to a coverage track on UCSC, but coverage is not visualized on the page itself.

There are a number of practical considerations for the optimal display of reference data. Specifically, as one primary use case for genome browsers involves gene-level analyses, the display of gene summary information is a central view for a genome browser, including integration of summary statistics as well as data for individual single nucleotide variants (SNV), insertions and deletions (indel), and copy number variants (CNV). Of course, detailed information on each variant, including annotations and quality metrics, is of paramount importance. However, an equally important display is that of the absence of variation: whether a missing variant implies a lack of observed variation, low or no coverage in the genomic region, or filtered variation.

The Exome Aggregation Consortium (ExAC) has collected, harmonized, and released exome sequence data from 60,706 individuals^3^. Already, these data have proven useful in filtering variants for identifying causal variants for rare disease^5–7^. Here, we present a visual browser of the ExAC dataset. The browser is intended for use by clinical geneticists researching variants of interest for patients as well as biologists exploring variation in specific genes.

## ExAC Browser

We designed the ExAC browser as an intuitive interface to enable clinical geneticists and biologists to explore variants and genes of interest. We built a scalable browser framework to display qualitative and quantitative information for genes and variants in the ExAC dataset (see Methods), including both quality control information as well as summary statistics. The front page of the browser includes a search bar, which is seeded with autocomplete suggestions based on gene symbols and aliases, as well as sample queries. From here, there are two central units of the ExAC browser: the gene (or transcript) page and the variant page.

### Gene/transcript page

The ExAC browser gene page is an overview page for gene-level information, including summary statistics, coverage, and variants. The page begins with gene metadata and external references, along with constraint information, which summarizes the gene's intolerance to variation for multiple functional classes ^3, 8^(Figure 1A). Next, we present single base-resolution coverage information for each exon for a number of metrics including mean, median, and proportion of individuals covered at a number of depth cutoffs (Figure 1B). Immediately below, an exon summary plot displays the position and frequency of each SNV and indel, as well as CNV count information broken down by population (Figure 1C). All individual CNV calls are provided in the form of UCSC tracks, which are linked at the top of the page and above the CNV display. Finally, the browser provides a comprehensive table for variant information, which includes the worst functional annotation across transcripts for each variant. The table is sortable, includes comprehensive frequency information, and can be exported to a CSV format (Figure 1D).

**Figure 1:**
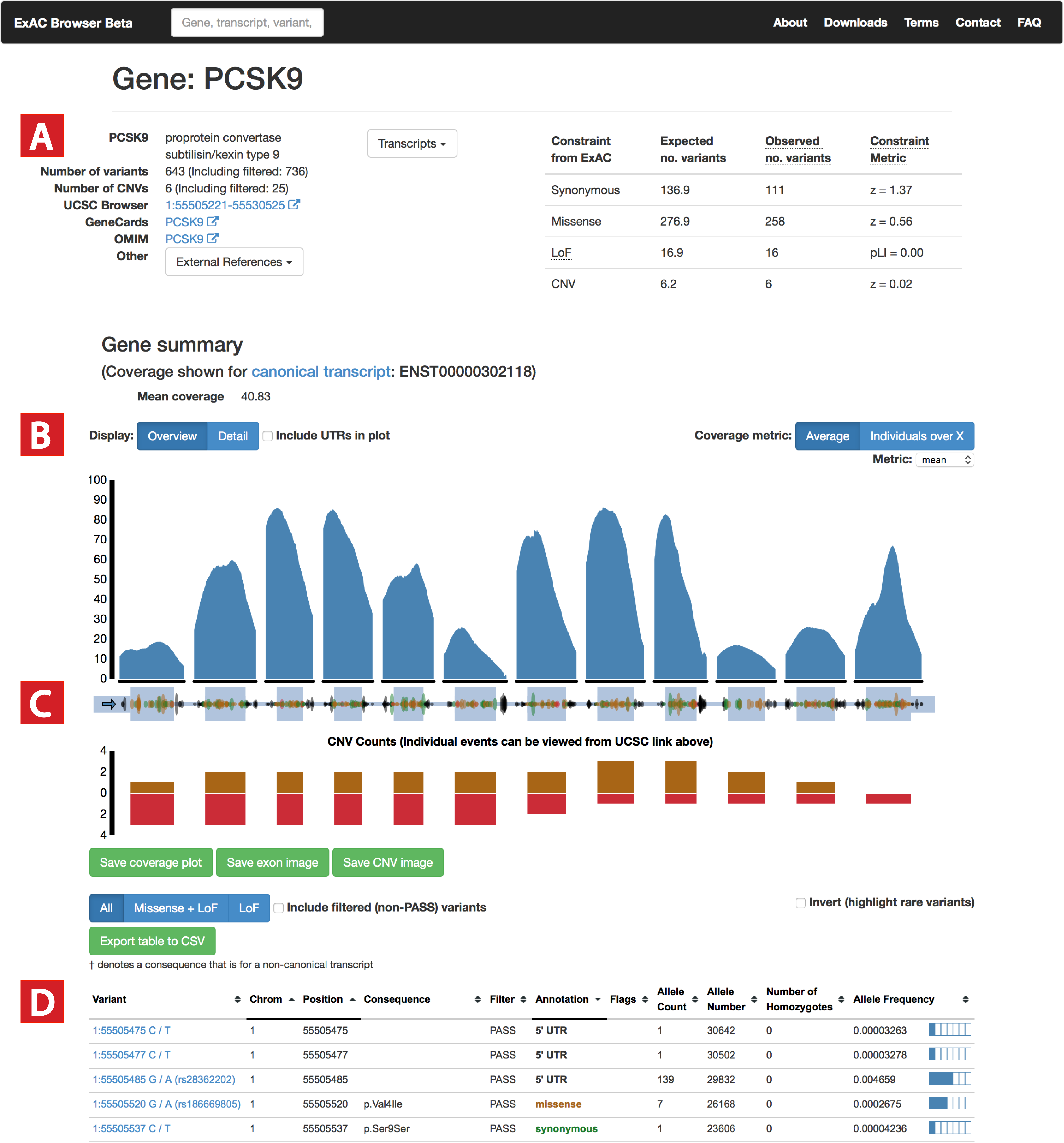
Gene page. (A): Gene information is summarized, including links to various external resources, as well as constraint information as described in^3^. For all exons in the canonical transcript, we display (B) base-level coverage for a number of metrics (mean coverage by default), as well as (C) position and frequency information for all variants, including CNVs. (D): A table of all variants is provided with additional annotation information and links to variant pages.

By default, the gene summary page presents a table of all variants in the gene, annotated with the worst consequence across all transcripts, as well as coverage information for the canonical transcript. We also present a transcript page that includes annotation and coverage information specific to that transcript.

### Variant page

The variant page includes a diverse set of annotations for the given variant. First, a site overview (Figure 2A) and site- and genotype-level quality metrics are provided (Figure 2B). The user is notified whether any individual has another variant in the same codon (suggesting a multi-nucleotide polymorphism, or MNP), whether the site is multi-allelic, or if a low number of individuals is covered at this locus. Functional annotations of the variant against each transcript including PolyPhen2^9^, SIFT^10^, and LOFTEE annotations, as well as a sortable table of population frequencies are provided (Figure 2C). Finally, for users that wish to evaluate the validity of specific variants, raw short-read data from a subset of individuals is available for each variant. We provide an IGVweb visualization of the read pileup of a 125 bp window around the variant (Figure 2D) for a random sample of individuals with each variant, as well as a sampling of homozygous individuals, if available. For the first time among genome browsers, we provide users with a mechanism to efficiently visualize the raw read support for a variant and make assessments of its quality that may not have been detected by variant calling algorithms.

**Figure 2:**
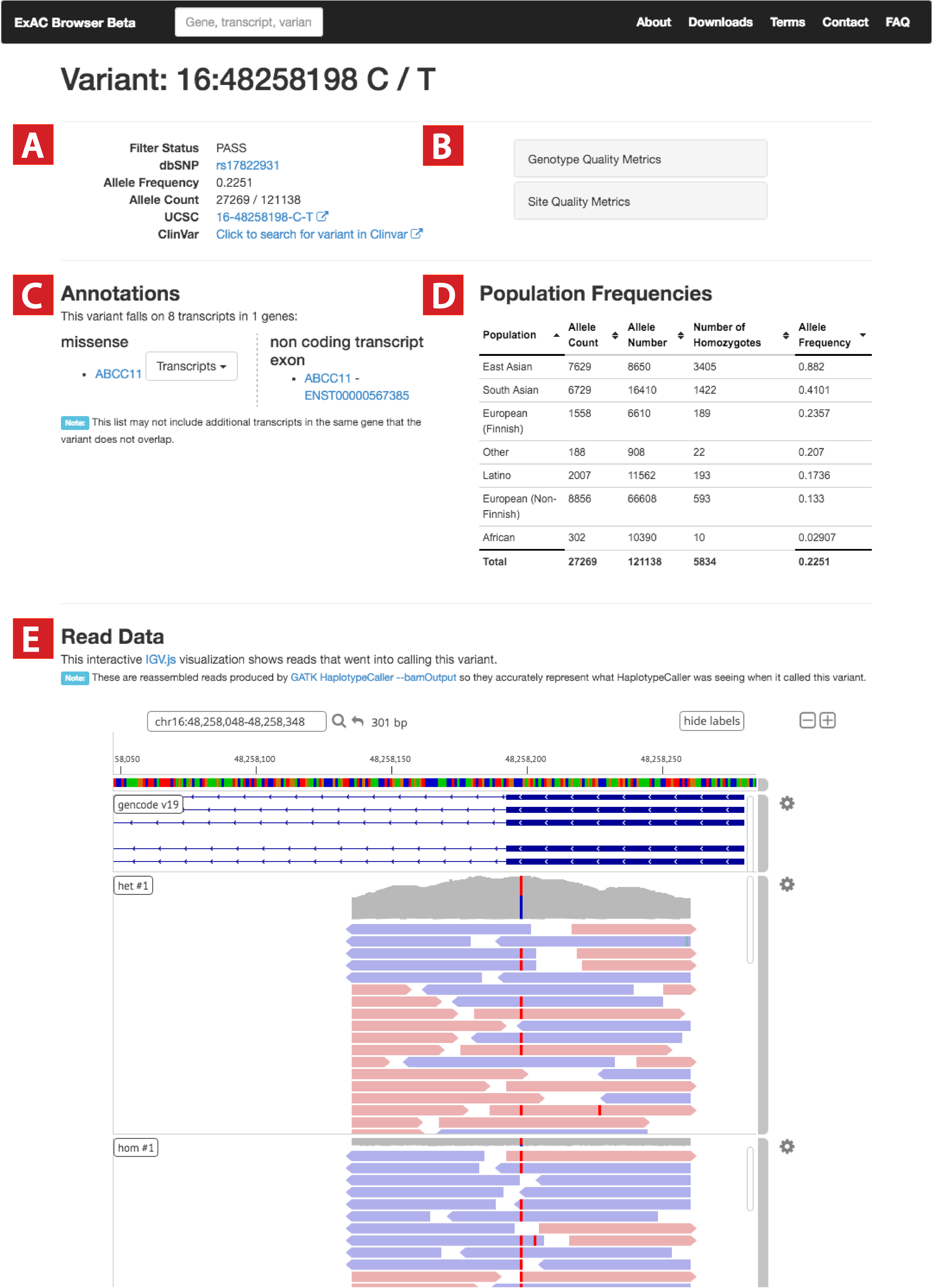
Variant page. (A): Variant metadata is displayed, including links to dbSNP, UCSC, and Clinvar. (B): Users can browse quality metrics based on genotypes (genotype quality and depth) as well as site-level quality metrics from GATK. (C): Annotations for each transcript are provided - if a variant overlaps multiple transcripts with the same functional annotation, a dropdown box provides additional details for the annotations. (D): Allele frequency information is displayed for each continental group (E): Short read data is provided for more technical users to assess validity of the variant call.

### Non-variant information

One important consideration for displaying genetic data includes the display of non-variant sites. In particular, if a variant or region is queried, we display metadata about the locus, whether or not variants are present in the dataset. When a user searches for variants or regions that are not covered in the ExAC dataset, the user is shown a page with coverage information for the general region for the variant.

### Additional considerations

As current web browsers and connections benefit from smaller data transfers and footprints, we have developed a number of optimizations to the browser, including compression and caching of data for large genes (Methods). Finally, the browser is optimized for mobile browsing, where extraneous information is hidden when browsing from a mobile device.

## Discussion

Here, we have described a browser for reference variation data, whose use has become widespread in clinical genetics laboratories across the world. As of this writing (8/1/2016), the browser has had over 250,000 users and 5 million pageviews spanning over 188 countries. The top 10 genes and top 3 variants visited by users are shown in Table 1.

**Table 1:** Top genes and variants viewed in the ExAC Browser

| Gene/Variant | Associated syndromes | Pageviews |
| --- | --- | --- |
| PCSK9 | Hypercholesterolemia (linked on front page of browser) | 13,540 |
| BRCA1 | Breast cancer susceptibility | 8,251 |
| BRCA2 | Breast cancer susceptibility | 7,408 |
| CFTR | Cystic Fibrosis | 5,179 |
| FBN1 | Marfan Syndrome | 4,886 |
| TP53 | Cancer susceptibility | 3,712 |
| TTN | Gene that encodes for the largest protein, cardiomyopathy | 3,528 |
| MYH7 | Cardiomyopathy | 3,497 |
| MYBPC3 | Cardiomyopathy | 3,398 |
| SCN5A | Brugada syndrome, long QT, cardiomyopathy | 3,175 |
| rs113993960 | Cystic fibrosis (CFTR deltaF508) | 203 |
| rs1799966 | Breast cancer (BRCA1 missense variant) | 157 |
| rs11571833 | Breast cancer (BRCA2 stop-gained variant) | 120 |

The code is open-source and available at http://github.com/konradjk/exac_browser. The browser framework established can be privately cloned and used for internal sequencing projects, as well as extended to a number of applications, such as a browser for results from genome-wide association studies (GWAS).

## Methods

### Data sources

As of this writing (8/1/2016), version 0.3.1 of ExAC dataset, as described in, was used for the ExAC Browser. Variants were annotated using the Variant Effect Predictor (VEP) version 81 ^11, 12^ against the Gencode v19 transcript set. dbSNP version 142 and dbNSFP was used for variant annotations, and gene names and aliases were extracted from dbNSFP ^13, 14^. Histograms for various genotype-specific quality metrics, such as per-sample genotype quality and depth, are pre-computed using a custom python script (https://github.com/macarthur-lab/exac_2015/blob/master/src/prepare_exac_sites_vcf.py). MNPs and constraint metrics are pre-calculated as described in^3^.

Reassembled read data was generated for each of the 9.8 million variants in ExAC v0.3.1 by running GATK HaplotypeCaller 3.1 (full version: v3.1-1-ga70dc6e) with the *-bamout* flag on each sample containing the particular variant (up to a limit of 5 homozygous and 5 heterozygous samples). Only samples with a read depth (DP) ≥ 10 and genotype quality (GQ) ≥ 20 were included. When a variant was present in more than 5 such samples, the 5 samples with the highest GQ were selected. Overall, HaplotypeCaller was run 22.3 million times to produce over 5Tb of small BAM files - with each BAM file storing reassembled reads for a several-hundred base pair window around the variant. Batches of several thousand of these small BAM files were then combined into larger BAM files to improve compression ratios, while using read groups to keep track of the original source of each read. The final dataset comprised ∼23,000 BAMs and spanned 540Gb. These BAM files were made directly available over the web and visualized in the ExAC browser using IGV.js.

Besides the *-bamout* flag, these additional flags were passed to HaplotypeCaller to ensure that gVCF genotype calls matched the original ExAC gVCF genotypes:

*-ERC GVCF --paddingAroundSNPs 300 --paddingAroundIndels 300 --max_alternate_alleles 3 -A DepthPerSampleHC -A StrandBiasBySample -- maxNumHaplotypesInPopulation 200 -stand_call_conf 30.0 -stand_emit_conf 30.0 -- disable_auto_index_creation_and_locking_when_reading_rods --minPruning 3 --variant_index_type LINEAR --variant_index_parameter 128000*

This data processing was managed by a python-based pipeline available here: https://github.com/macarthur-lab/exac_readviz_scripts CNVs were generated using XHMM^15^ and based on GENCODE v19 coding regions: all details of CNV calling and quality control have been published previously^16^. Gene summary CNV counts and related constraint scores are presented based on likelihoods of the CNV occurring within the genomic range of the gene, as described^16^. Exon CNV counts and CNVs presented in the UCSC browser are based on all confidently called CNVs (XHMM SQ > 60) across the genome. All overlapping CNVs, regardless of amount of overlap, are included in Exon CNV counts.

### Website design

The ExAC browser was built primarily on open-source tools. On the server, a lightweight Flask framework serves content built on Python scripts available at http://github.com/konradjk/exac_browser. All variants and metadata are loaded into MongoDB (version 2.4.14). The major components loaded include the variant data (directly from the VCF format), coverage data (generated by a modified version of samtools, as described in^3^), MNP and constraint information, as well as gene models from Gencode and RSID information from dbSNP.

The HTML backbone was created based on Bootstrap version 3.1.1 (https://github.com/twbs/bootstrap) and JQuery version 1.11.1 (http://jquery.org). Plotting was performed using d3 version 3 ^17^. Read visualization is powered by IGVweb version 0.9.3 (https://github.com/igvteam/igv.js/releases/tag/0.9.3).

The entire system runs on a Linux virtual machine with 8 cores, 32 GB RAM, and 2 T of disk space using Apache 2.4.12. Page tracking is provided by Google Analytics (http://www.google.com/analytics/).

### Optimizations

Bootstrap is a mobile-first web framework, which enables the ExAC browser's optimizations for mobile browsing: specifically, much extraneous information (such as the coverage information or additional variant annotations) is hidden when the browser is used on a smaller screen. Additionally, the pages for large genes are pre-computed, allowing for faster load times for these genes. Finally, user search is optimized using typeahead version 0.10.2, with most search terms, including gene names and all aliases, populating the search bar. The single search bar is used to search for variants (formatted as RSIDs or in a chromosome and position format), genes and transcripts (symbols, aliases, or Ensembl identifiers), and regions.

## Acknowledgments

K.J.K. is supported by NIGMS Fellowship (F32GM115208). M.L. is supported by the Australian National Health and Medical Research Council CJ Martin Fellowship, Australian American Association Sir Keith Murdoch Fellowship and the MDA/AANEM Development Grant. D.G.M is supported by NIGMS R01 GM104371 and NIDDK U54 DK105566.

## References

1. Consortium, 1. G. P. An integrated map of genetic variation from 1,092 human genomes. Nature (2012). doi:doi:10.1038/nature11632

2. Tennessen, J. A. et al. Evolution and functional impact of rare coding variation from deep sequencing of human exomes. Science 337, 64–69 (2012).

3. Lek, M. et al. Analysis of protein-coding genetic variation in 60,706 humans. Nature 536, 285–291 (2016).

4. Kent, W. J. et al. The human genome browser at UCSC. Genome Res 12, 996–1006 (2002).

5. Grozeva, D. et al. Targeted Next-Generation Sequencing Analysis of 1,000 Individuals with Intellectual Disability. Hum Mutat 36, 1197–1204 (2015).

6. Robinson, E. B. et al. Genetic risk for autism spectrum disorders and neuropsychiatric variation in the general population. Nat Genet 48, 552–555 (2016).

7. Song, W. et al. Exploring the landscape of pathogenic genetic variation in the ExAC population database: insights of relevance to variant classification. Genet Med (2015). doi:10.1038/gim.2015.180

8. Samocha, K. E. et al. A framework for the interpretation of de novo mutation in human disease. Nat Genet (2014). doi:10.1038/ng.3050

9. Adzhubei, I. A. et al. A method and server for predicting damaging missense mutations. Nature Methods 7, 248–249 (2010).

10. Kumar, P., Henikoff, S. & Ng, P. C. Predicting the effects of coding non-synonymous variants on protein function using the SIFT algorithm. Nat Protoc 4, 1073–1081 (2009).

11. McLaren, W. et al. Deriving the consequences of genomic variants with the Ensembl API and SNP Effect Predictor. Bioinformatics 26, 2069–2070 (2010).

12. McLaren, W. et al. The Ensembl Variant Effect Predictor. Genome Biol 17, 1 (2016).

13. Liu, X., Jian, X. & Boerwinkle, E. dbNSFP: a lightweight database of human nonsynonymous SNPs and their functional predictions. Hum Mutat 32, 894–899 (2011).

14. Liu, X., Jian, X. & Boerwinkle, E. dbNSFP v2.0: A Database of Human Non-synonymous SNVs and Their Functional Predictions and Annotations. Hum Mutat 34, E2393–E2402 (2013).

15. Fromer, M. et al. Discovery and Statistical Genotyping of Copy-Number Variation from Whole-Exome Sequencing Depth. The American Journal of Human Genetics 91, 597–607 (2012).

16. Ruderfer, D. M. et al. Patterns of genic intolerance of rare copy number variation in 59,898 human exomes. Nat Genet (2016). doi:10.1038/ng.3638

17. Bostock, M. D3. js Overview: D3 Data-Driven Documents. (2015).

